# ANARCII: A Generalised Language Model for Antigen Receptor Numbering

**DOI:** 10.1101/2025.04.16.648720

**Authors:** Alexander Greenshields-Watson, Parth Agarwal, Sarah A. Robinson, Benjamin Heathcote Williams, Gemma L. Gordon, Henriette L. Capel, Yushi Li, Fabian C. Spoendlin, Broncio Aguilar-Sanjuan, Fergus Boyles, Charlotte M. Deane

**Affiliations:** Oxford Protein Informatics Group, Department of Statistics, University of Oxford, 24-29 St Giles’, Oxford, OX1 3LB United Kingdom

## Abstract

Antigen receptor numbering allows the rapid delineation of the antigen-binding regions of antibody and T cell receptor (TCR) sequences, from sequence alone. It also allows the comparison of the vast diversity of antigen receptors in a consistent frame of reference. Numbering of antigen receptors is currently achieved by aligning sequences to a reference set. This approach may result in different numbering, depending on the reference set used or may fail to number query sequences derived from new species or rare sequence types. To address this problem, we have built a new numbering method (ANARCII) which requires no alignment step and is based on a Seq2Seq language model.

Our results show that ANARCII can deal with the complexity that arises in experimentally collected sequencing data and generalise to sequences which are highly dissimilar to those in training. In test sets designed to contain challenging and ambiguous sequence patterns ANARCII numbering was identical to existing methods for over 99.99% of conserved residues and over 99.94% for complete CDR regions. The lightweight architecture allows numbering of over 90,000 sequences per minute on a single A100 GPU. Furthermore, the ANARCII package can be conditioned to fit rare sequence types and provide new training data for fine-tuning. We demonstrate that fine-tuned versions of ANARCII can correctly number other immunoglobulin domains such as TCRs and VNARs. Our model is freely available as a web tool (https://github.com/oxpig/ANARCII), as well as a package for high throughput numbering of next generation sequencing data (https://opig.stats.ox.ac.uk/webapps/sabdab-sabpred/sabpred/anarcii/).

## INTRODUCTION

Standard antibodies are Y-shaped proteins made up of two pairs of identical heavy and light chains. Their ability to bind to target epitopes with high affinity and specificity makes them highly attractive therapeutics (Carter and Rajpal 2022). The antibody structure can be divided into the constant region, formed of the lower two domains of each heavy chain, and the fragment antigen binding (Fab), made up of the entire light chain and upper two domains of the heavy chain. The distal domains of the Fab (VH and VL) form the variable region (Fv) on which are found the complementarity determining region loops (CDR1, CDR2 and CDR3), the main segments of an antibody involved in antigen binding.

In antibody heavy chains, sequence diversity arises from the assembly of variable (V), diversity (D) and joining (J) gene segments to form a complete chain (Brack et al. 1978), in a process termed VDJ recombination. The genome of an individual contains multiple distinct V, D and J genes; the steps and enzymes involved in joining these genes introduce random mutations giving rise to junctional diversity (Alt et al. 1984). The junctions are situated in the CDR3 region of the heavy chain (CDRH3), leading to this loop being the focus of maximal sequence diversity and a primary driver of antigen recognition (Regep et al. 2017). Additionally, when an antibody undergoes affinity maturation to a target antigen, the process of somatic hypermutation (SHM) introduces further random mutations along the sequence (Papavasiliou and Schatz 2002). Mutations which result in higher affinity for the target are positively selected and may dominate the immune response. In light chains the process is identical, except for the absence of a D gene in the generation process.

Application of numbering schemes to the antibody Fv region provides a way to navigate the vast diversity that these mechanisms generate and compare many sequences using a consistent frame of reference. The substitutions, insertions and deletions that arise in SHM can be identified and numbered with respect to the original parent germline gene sequence. The primary purpose of numbering an Fv region is to delineate the start and end of the CDR loops from the beta sheet framework regions (FR) from which they protrude (Dondelinger et al. 2018). There are several different schemes which are used to number antibodies. The earliest suggested scheme is the Kabat scheme (Wu and Kabat 1970). This was created without any structural information and using only a small number of sequences. More structurally aware derivates of the Kabat scheme, Chothia (Chothia and Lesk 1987) and Martin (Abhinandan and Martin 2008) followed. Further developments included the Aho scheme (Honegger and Plückthun 2001) which sought to be highly representative of structure, and the Wolfguy system (Bujotzek et al. 2015) which allowed compliance with docking tools that do not accept repeated use of the same number (a problem when docking heavy and light chains). The most widely used numbering scheme is IMGT (Lefranc et al. 2015). First designed in 1997 (V. Giudicelli et al. 1997), the scheme has consistent placement of conserved residues and regions across antibody heavy and light chains, as well as TCR alpha, beta, gamma and delta chains. As it utilises the germline reference sequences as the basis for numbering, insertions and deletions can be identified with respect to the parent allele. It is conceptually simple to understand, and the structural equivalence of numbering positions makes it easy to compare across sequence types, chains, genes and alleles.

Currently, antibodies are numbered by aligning the sequence of an antibody to either a consensus sequence or the germline reference sequences. Tools including AbRSA (L. Li et al. 2019) and AntPack (Parkinson and Wang 2024) align to a consensus sequence using BLOSUM62 (Henikoff and Henikoff 1992), custom gap penalties and the Needleman-Wuncsh algorithm (Needleman and Wunsch 1970). IMGT V-quest (Véronique Giudicelli, Chaume, and Lefranc 2004) and IgBLAST (Ye et al. 2013) use Smith-Waterman (Smith and Waterman 1981) or BLAST (Altschul et al. 1990), respectively to align against IMGT reference sequences and return the highest scoring alignment. An alternative approach is used by ANARCI (pronounced ‘anarchy’) (Dunbar and Deane 2016), which scores a query sequence against multiple Hidden Markov Models (HMMs) built on sets of germline sequences for a given species and chain.

All of these methods rely on either a representative consensus sequence or an IMGT germline reference set compiled using data available at a specific point in time. Germline references are continually being updated and modified as new species, genes, alleles and sequence types are discovered (Collins et al. 2024). This means the ground truth needed to number an antibody is not static, and the numbering returned may differ depending on the timestamp of the reference set. For example, users of ANARCI have identified differences in numbering depending on when the HMM models were compiled (see GitHub issues #40 and #64 at https://github.com/oxpig/ANARCI). These differences may occur in up to 4% of sequences taken from random samples of Observed Antibody Space (Olsen, Boyles, and Deane 2022; Kovaltsuk et al. 2018), and mainly occur in the starting residues of truncated fragments, as well as the CDRH3.

Furthermore, query sequences which originate from rare species or uncommon sequence formats (e.g. novel single chain constructs) are more challenging to number with current approaches which rely on static sequence representations. This is because the sequences can be very different from the data used to construct the consensus sequences or HMMs, and the existing methods cannot generalise to novel sequence types. This problem is becoming increasingly relevant as more antibody sequence data from new species, alleles and sequence types is deposited in the public domain. One example of this are shark variable domain of new antigen receptor (VNAR) antibodies (Simmons et al. 2006), where the large numbers of proximal cysteines, as well as non-standard CDRH2 loop placement, mean that existing tools cannot provide numbering appropriate for their unique structural morphology (Kovalenko et al. 2013).

Large language models (LLMs) have been successful in many protein sequence/structure tasks (Lin et al. 2023; Hie et al. 2023) suggesting that they may capture the sequence patterns which relate to structure. In this paper we explore whether a language model (Vaswani et al. 2017) can accurately number antigen receptors and effectively generalise to new sequence types and patterns.

Models were trained on over 160M antibody heavy and light chain sequences. Each sequence was augmented with random, non-antibody sequence content. During training models were tasked with distinguishing antibody sequence content from non-antibody content, predicting the correct chain identity (heavy, kappa or lambda) and applying IMGT numbering to the sequence. Multiple model architectures were investigated to find the optimum balance of inference speed and numbering accuracy. The final model suite, ANARCII (pronounced “anarchy two’’), was able to replicate the results of ANARCI, whilst generalising to sequence types which could not previously be numbered in any form.

ANARCII outputs can also be conditioned to adapt predicted numbering to highly novel sequence types. We exploited this to number 456 shark VNAR sequences with unconventional sequence features so that they fully matched the IMGT definition around the CDR2. A version of the ANARCII model fine-tuned on this renumbered VNAR dataset could accurately predict the correct numbering on held-out VNAR test sequences without conditioning. We also demonstrate that the ANARCII antibody models with only minimal fine-tuning can accurately number T cell receptor (TCR) sequences. Multiple model types are made accessible to users in the ANARCII software suite, as well as instructions on how to condition auto-regressive outputs to create custom schemes and workflows. ANARCII will be increasingly useful to antibody researchers and engineers as they navigate the increasingly vast landscape of antibody sequence space. It also serves as a proof of concept that small language models can be effective in sequence alignment tasks, offering a flexible alternative to current methods.

## RESULTS

### Training a Seq2Seq Language model to number antigen receptor sequences

In order to develop a model that could “translate” the amino acid sequences of antigen receptors into their corresponding IMGT number labels we utilised the Seq2Seq transformer architecture and the neural machine translation task. Our model, ANARCII, is designed to both number and call the chain (heavy, kappa or lambda) of a given antibody Fv sequence. Figure 1A shows a schematic of the training task, in which the model learns to predict [<CHAIN>], numbering (IMGT numbers 1-128) and insertion [<X>] tokens from an input amino acid sequence (**Figure 1A**). Sequences found in the PDB (Berman et al. 2000) or in experimental data (Olsen, Boyles, and Deane 2022) often contain extra content such as leader sequences. To ensure that ANARCII can correctly number experimental data of this type, we randomly added between 0-40 amino acids to the start and end of every sequence during training (termed “junk” in **Figure 1A**) and trained the model to output a [<SKIP>] token as the prediction, with the [<EOS>] token directly after the antibody sequence content.

**Figure 1.**
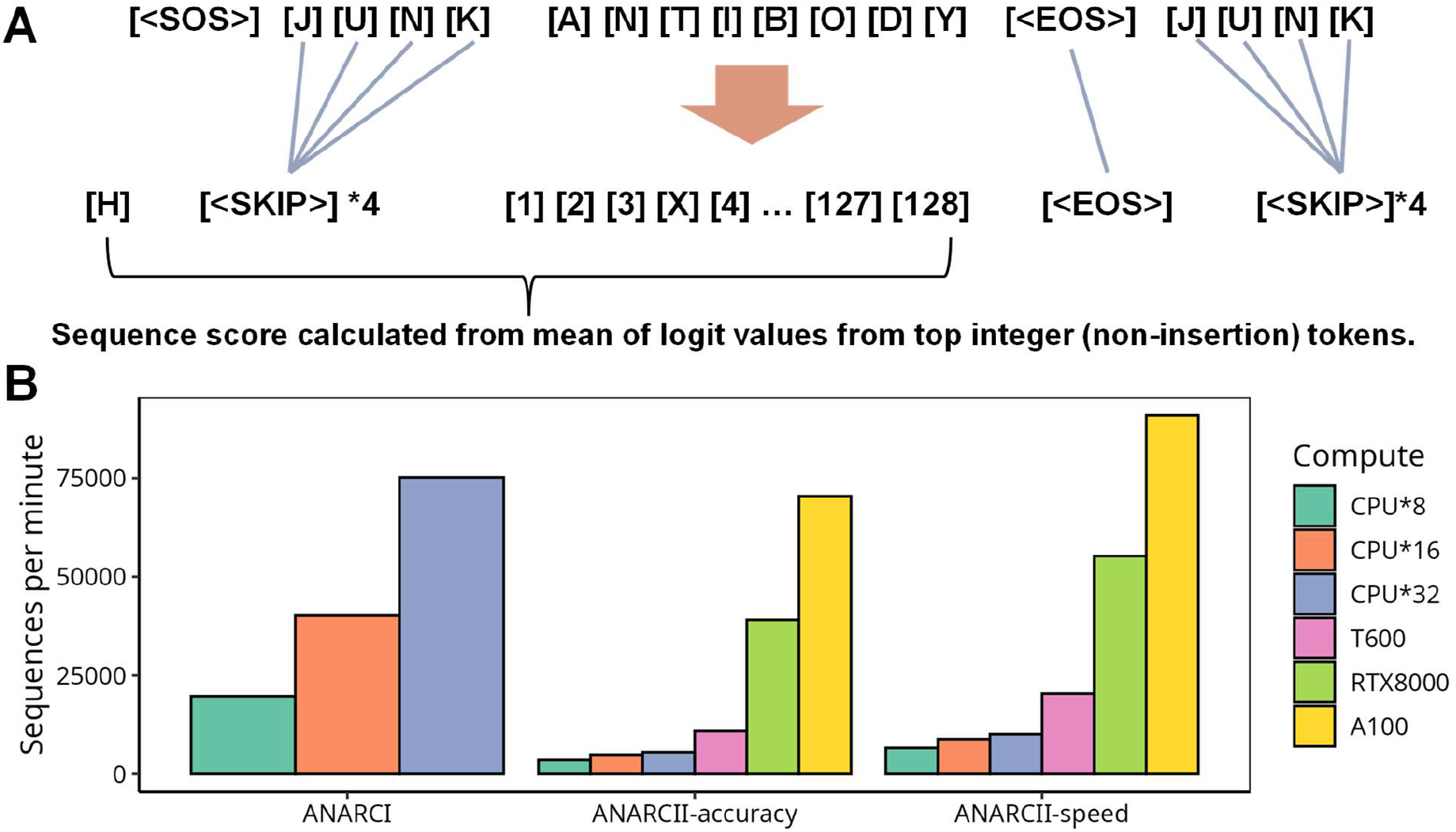
ANARCII is a Seq2Seq language model trained to identify chain types and number sequences. Schematic of sequence to number training task (**A**), where input sequences are bookended by random amino acids, which during training must be assigned the corresponding [<SKIP>] token. In addition, each target sequence has a chain token which the model is trained to predict before assigning numbering. (**B**) Comparison of inference speeds on 100,000 sequences across different CPU and GPU architectures for ANARCI, and the ANARCII models.

As described in the methods, the training data was ∼95.7M heavy, ∼41.6M kappa and ∼33.4M lambda non-redundant sequences sampled from the Observed Antibody Space (OAS). Heavy, kappa and lambda sequences were each clustered based on sequence identity, then the resulting clusters split into train, test and validation sets (90:5:5, see methods). Hyperparameters for ANARCII were chosen based on the validation set accuracy and the corresponding inference speed (**Supp Table 2**). While large parameter models show the best validation loss, often the difference in numbering accuracy was very small (differing between 0.1-0.2%) and did not justify the increases in inference time associated with having more parameters. We created two versions of ANARCII: *ANARCII-accuracy* and *ANARCII-speed*. The original ANARCI (without calling V/J gene usage) can number ∼75,000 sequences per minute (SPM) using 32 CPUs. *ANARCII-accuracy* on an A100 GPU can number ∼70,000 SPM, while *ANARCII-speed* can number ∼90,000 SPM on the same architecture (**Figure 1B**).

**Table 1.** ANARCII models performance on held out test sequences (including truncated sequences). % Agreement with ANARCI.

| Test Set | Model | Overall | CYS-23 | CYS-104 | TRP-41 | Ar-118 | CDR1 | CDR2 | CDR3 | DE-Loop |
| --- | --- | --- | --- | --- | --- | --- | --- | --- | --- | --- |
| Heavy | Speed | 99.8233 | 99.9999 | 99.9992 | 99.9999 | 99.9969 | 99.9898 | 99.9578 | 99.9747 | 99.9826 |
| Heavy comp |  | 99.9545 | 99.9999 | 99.9994 | 99.9999 | 99.9971 | 99.9947 | 99.9619 | 99.9895 | 99.9839 |
| Heavy junk |  | 96.095 | 99.9986 | 99.9964 | 99.9908 | 99.9943 | 99.8105 | 99.6688 | 99.7777 | 99.9126 |
| Kappa |  | 99.5937 | 99.9982 | 99.9936 | 99.9974 | 99.9983 | 99.7911 | 99.8373 | 99.9321 | 99.7121 |
| Kappa comp |  | 99.8393 | 99.9988 | 99.9959 | 99.9984 | 99.9991 | 99.9629 | 99.8866 | 99.9849 | 99.7585 |
| Kappa junk |  | 96.0655 | 99.9961 | 99.9931 | 99.9972 | 99.9992 | 99.4871 | 99.8209 | 99.8755 | 99.7636 |
| Lambda |  | 99.5378 | 99.9966 | 99.9927 | 99.9986 | 99.998 | 99.7235 | 99.8559 | 99.9348 | 99.8478 |
| Lambda comp |  | 99.8652 | 99.9971 | 99.9943 | 99.9987 | 99.9983 | 99.9648 | 99.8958 | 99.978 | 99.8645 |
| Lambda junk |  | 97.3685 | 99.9925 | 99.9891 | 99.9984 | 99.9974 | 99.2406 | 99.6707 | 99.8151 | 99.7704 |
| Heavy | Accuracy | 99.8660 | 99.9999 | 99.9998 | 99.9999 | 99.9985 | 99.9928 | 99.9732 | 99.9832 | 99.9925 |
| Heavy comp |  | 99.9744 | 100 | 99.9998 | 99.9999 | 99.9985 | 99.9978 | 99.9771 | 99.995 | 99.9943 |
| Heavy junk |  | 96.4897 | 99.9995 | 99.9987 | 99.9997 | 99.9974 | 99.8687 | 99.8316 | 99.8235 | 99.9539 |
| Kappa |  | 99.6746 | 99.999 | 99.998 | 99.9979 | 99.9995 | 99.8210 | 99.8919 | 99.9606 | 99.8314 |
| Kappa comp |  | 99.897 | 99.9991 | 99.9985 | 99.9987 | 99.9997 | 99.9862 | 99.9372 | 99.9948 | 99.8682 |
| Kappa junk |  | 96.5836 | 99.9983 | 99.9974 | 99.9991 | 99.9998 | 99.5863 | 99.875 | 99.9139 | 99.8404 |
| Lambda |  | 99.6249 | 99.998 | 99.997 | 99.9996 | 99.9985 | 99.7544 | 99.9028 | 99.9539 | 99.9078 |
| Lambda comp |  | 99.9047 | 99.9982 | 99.9975 | 99.9997 | 99.9987 | 99.9751 | 99.9408 | 99.9887 | 99.9179 |
| Lambda junk |  | 97.7400 | 99.9974 | 99.9943 | 99.999 | 99.9994 | 99.3573 | 99.7003 | 99.8697 | 99.8458 |

**Table 2.** ANARCII models performance on full length non-truncated test sequences. % Agreement with ANARCI.

| Test set | Model | Overall | CYS-23 | CYS-104 | TRP-41 | Ar-118 | CDR1 | CDR2 | CDR3 | DE-Loop |
| --- | --- | --- | --- | --- | --- | --- | --- | --- | --- | --- |
| Heavy | Speed | 99.8192 | 99.9999 | 99.9998 | 99.9999 | 99.9992 | 99.9963 | 99.9838 | 99.9953 | 99.9907 |
| Heavy comp |  | 99.9544 | 99.9999 | 99.9999 | 99.9999 | 99.9993 | 99.9969 | 99.9834 | 99.9974 | 99.9909 |
| Heavy junk |  | 95.9818 | 99.9988 | 99.9988 | 99.9991 | 99.999 | 99.979 | 99.8581 | 99.9808 | 99.945 |
| Kappa |  | 99.5819 | 99.9998 | 99.9994 | 99.9998 | 100 | 99.996 | 99.9695 | 99.9965 | 99.9361 |
| Kappa comp |  | 99.8389 | 99.9998 | 99.9994 | 99.9998 | 100 | 99.9976 | 99.9669 | 99.9982 | 99.9340 |
| Kappa junk |  | 95.8878 | 99.9996 | 99.9991 | 99.9991 | 99.9998 | 99.9879 | 99.9805 | 99.9928 | 99.9560 |
| Lambda |  | 99.5134 | 99.9998 | 99.9981 | 99.9996 | 99.9999 | 99.9906 | 99.9454 | 99.9878 | 99.9092 |
| Lambda comp |  | 99.8648 | 99.9999 | 99.9981 | 99.9996 | 99.9999 | 99.9984 | 99.9458 | 99.994 | 99.9116 |
| Lambda junk |  | 97.2163 | 99.9995 | 99.998 | 99.9995 | 99.9994 | 99.9842 | 99.9495 | 99.9776 | 99.8939 |
| Heavy | Accuracy | 99.8632 | 100 | 100 | 100 | 99.9996 | 99.9987 | 99.9922 | 99.9969 | 99.9962 |
| Heavy comp |  | 99.9744 | 100 | 100 | 100 | 99.9996 | 99.9991 | 99.9919 | 99.9987 | 99.9963 |
| Heavy junk |  | 96.4010 | 99.9996 | 99.9996 | 99.9998 | 99.9996 | 99.9887 | 99.9549 | 99.9875 | 99.9732 |
| Kappa |  | 99.6645 | 99.9999 | 99.9999 | 99.9998 | 99.9999 | 99.9963 | 99.9902 | 99.9936 | 99.9823 |
| Kappa comp |  | 99.8968 | 99.9998 | 99.9998 | 99.9998 | 99.9998 | 99.9979 | 99.9905 | 99.9991 | 99.9846 |
| Kappa junk |  | 96.4539 | 99.9999 | 99.9998 | 99.9997 | 100 | 99.9953 | 99.9889 | 99.9975 | 99.9743 |
| Lambda |  | 99.6068 | 99.9998 | 99.9994 | 99.9998 | 100 | 99.9917 | 99.9665 | 99.9934 | 99.9457 |
| Lambda comp |  | 99.9045 | 99.9999 | 99.9995 | 99.9998 | 100 | 99.9986 | 99.9684 | 99.9984 | 99.9494 |
| Lambda junk |  | 97.6322 | 99.9997 | 99.9988 | 99.9997 | 99.9998 | 99.9896 | 99.9725 | 99.988 | 99.9357 |

### ANARCII models can consistently and accurately IMGT number antigen receptor sequences

To investigate the accuracy of our ANARCII models we used them to number the held-out test sets of ∼9.6M heavy, ∼1.5M kappa and ∼2.4M lambda sequences and compared the results against ANARCI. These test sets included sequences of varying length with many start or end truncations typically observed in NGS datasets. *ANARCII-speed* exactly replicated the ANARCI numbering on 99.82% (heavy), 99.59% (kappa) and 99.54% (lambda) of the respective test sets (**Table 1**), while *ANARCII-accuracy* agreed on 99.87%, 99.67% and 99.62%. We next explored where the numbering predictions diverged between ANARCI and ANARCII and found that most disagreements were at the ends of truncated test sequences where identification of the first or last residues were not clear, and the remainder of the sequence numbering was identical (far left region of plots in **Figure 2A-C**). We also found that most divergence, especially in light chains, occurred in sequences which did not contain conserved residues such as Cys104 or Trp41.

**Figure 2.**
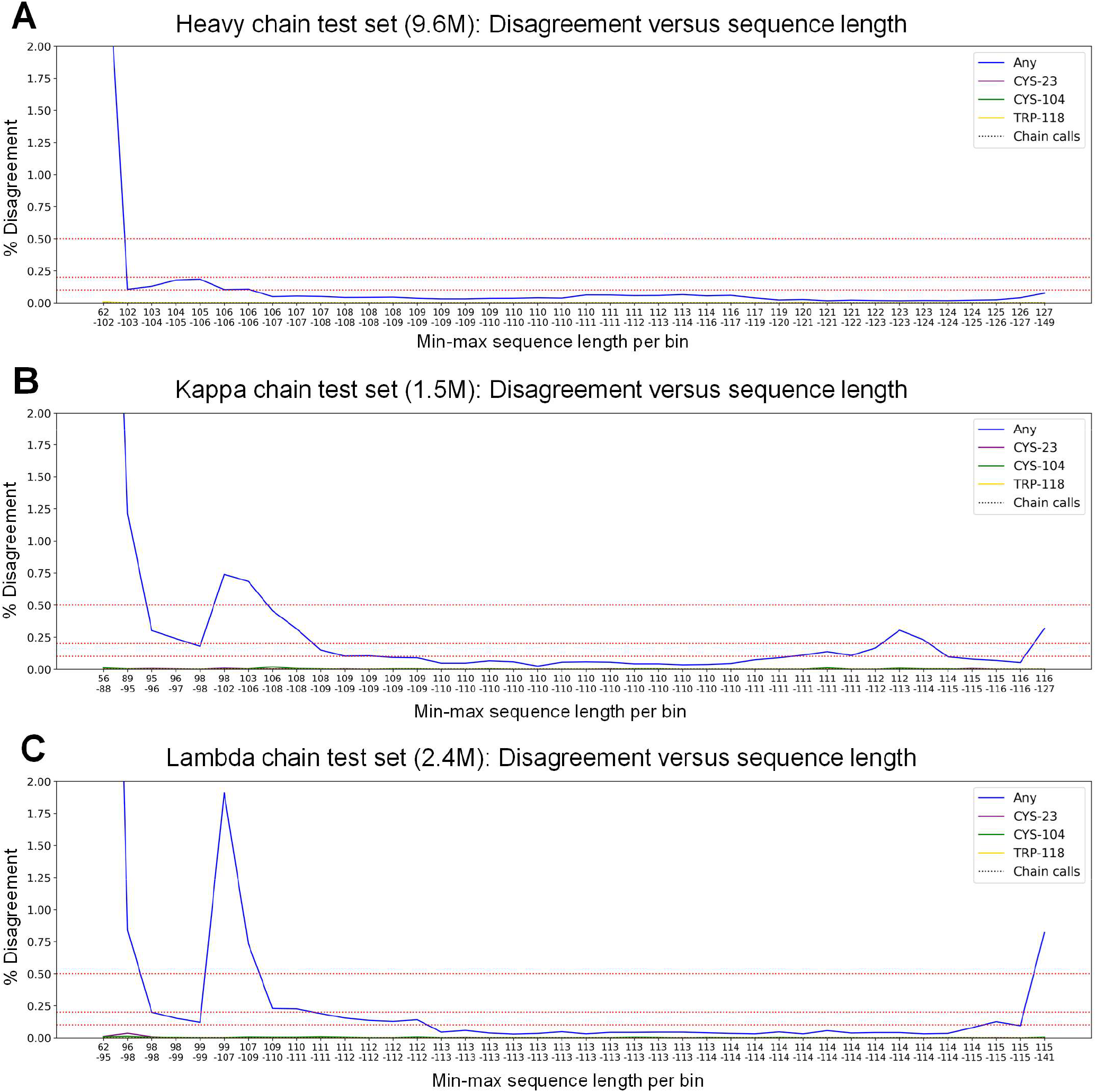
Sequence with truncations are the main source of disagreement between HMM (ANARCI) and LM (ANARCII-accuracy) numbering. Disagreement in numbering of ANARCI and ANARCII-accuracy versus sequence length on held out test sets of heavy (**A**), kappa (**B**) and lambda (**C**) chains. Test sequences are divided into 40 bins based on length, indicated on the x axes, from short to longer sequences.

*ANARCII-accuracy* was able to correctly number the conserved cysteines (23, 104) and aromatics (41 and 118) residues with over 99.99% agreement to ANARCI in both heavy, kappa and lambda chains (**Table 1**). Agreement of all the positions of CDR regions was lower, with a minimal agreement of 99.97% for heavy, 99.82% for kappa and 99.75% for lambda chains. By filtering the test sets to ensure all target sequences contained conserved cysteines at positions 23 and 104 we increased the CDR delineation agreement to above 99.98%, 99.94% and 99.94% in heavy, kappa and lambda chains (**Table 1** – row labelled ‘comp’). The addition of random sequence (termed junk) negatively impacted overall agreement (**Table 1** – row labelled ‘junk ends’), however this was also related to truncations where assignment of the start or end residue was further complicated by the presence of random residues (some of which may have resembled viable antibody content). In the CDR regions the agreement values were slightly lower for *ANARCII-speed*, however most values were reduced by less than 0.1% on these challenging test sequences (**Table 1**).

Given that truncations had a negative impact on agreement we filtered our test sets to remove these sequences and recalculated agreement with ANARCI numbering (**Table 2**). This boosted CDR1 and CDR3 agreement to 99.99% for all chains using *ANARCII-accuracy*, while conserved residue agreement was close to 100% (lowest value of 99.9988%, lambda chains with junk). CDR2 agreement was lowest at 99.95% for heavy chains, 99.99% for kappa and 99.97% for lambda chains. Truncated sequences are commonly found in NGS datasets (Olsen, Boyles, and Deane 2022) and the absence of start or end residues creates challenges for both ANARCI (HMM) and ANARCII (language models) and could explain the poor agreement in this area. However, this could not explain the disagreement within the CDR2 and DE-loop (Kelow, Adolf-Bryfogle, and Dunbrack 2020) regions where agreement was worst despite this region being less diverse than the CDR3 region. We next investigated the numbering of sequences which were inconsistently labelled between different versions of ANARCI to see whether the CDR2 and DE-loop region was also the primary source of ambiguity and whether the use of an additional numbering method could help find a ground truth.

### Numbering of challenging sequences highlights limitations of alignment methods and reflects ambiguity in the CDR2 to DE-loop region

As described in the methods we identified sequences where the number labels assigned by the latest version of ANARCI diverged from the initial numbering deposited in OAS. These differences could be explained by changes in the germline reference sequences used to construct the HMMs which underpin ANARCI. For example, the addition of new alleles to the IMGT reference set may result in alternate numbering of certain sequences between versions of ANARCI built with and without this updated information. We filtered these sequences to identify those with the greatest difference between ANARCI versions to create a challenging dataset, the ‘*no-truth*’ set. We numbered the *no-truth* set sequences with ANARCI, *ANARCII-accuracy, ANARCII-speed* and a rapid alignment-based tool, AntPack (Parkinson and Wang 2024).

Analysis of numberings in the *no-truth* set revealed that agreement was highest between *ANARCII-speed* and *ANARCII-accuracy*, with 91.1% of sequences numbered identically, and most disagreements falling in the CDR3 region (**Supp Fig. 2A-B**). As *ANARCII-accuracy* showed marginally higher identification of conserved Cys104 residues (276 more identified, difference of <0.1% in across all ∼900K no-truth sequences) we focused on comparisons between *ANARCII-accuracy*, ANARCI and AntPack for the remainder of the analysis.

When compared to ANARCI (recent version), *ANARCII-accuracy* (**Figure 3A**) numbered 52.4% of heavy sequences identically. In sequences where numberings were not identical, labels mainly diverged in the CDRH3 in stretches of sequence containing multiple aromatic, or non-standard residues around position 118 (for example runs of repeated tyrosine residues from positions 116 to 119). Comparison of numberings between AntPack with *ANARCI (49*.*5% of sequences with identical numbering)* and AntPack with *ANARCII-accuracy (54*.*9% identical)* were similar, however AntPack disagreed with *ANARCI* and *ANARCII-accuracy* at Cys23 in 0.7 % of sequences and Cys104 in 4.0% of sequences. Inspection of these disagreements revealed that AntPack was indicating a non-Cys amino acid at positions-23 and 104, in 0.5% and 4.0% of *no-truth* heavy sequences respectively. Of the three methods *ANARCII-accuracy* mislabelled the fewest Cys23 and Cys104 residues, with values of less than 0.1%. Further analysis showed most of the affected sequences had a truncation in the first 50 residues. If the truncation occurred before the Cys23, AntPack would begin with a number label close to 1, wrongly assigning the conserved cysteine then “catching up” by placing a large ∼10 residue deletion in the CDR1 so that the subsequent sequence was correct (correct identification of Trp41) (**Supp Fig. 2C**). If the sequence was truncated at the CDR2 region (residues 50-60), then AntPack would again number with a label between 1-20, incorrectly assigning the entire sequence and placing a large deletion in the CDR3 to allow correct numbering of the aromatic at position 118. We also noted that several sequences where *ANARCI* and *ANARCII-accuracy* agreed, but AntPack did not, contained ‘X’ at positions early in the sequence, in place of standard amino acids.

**Figure 3.**
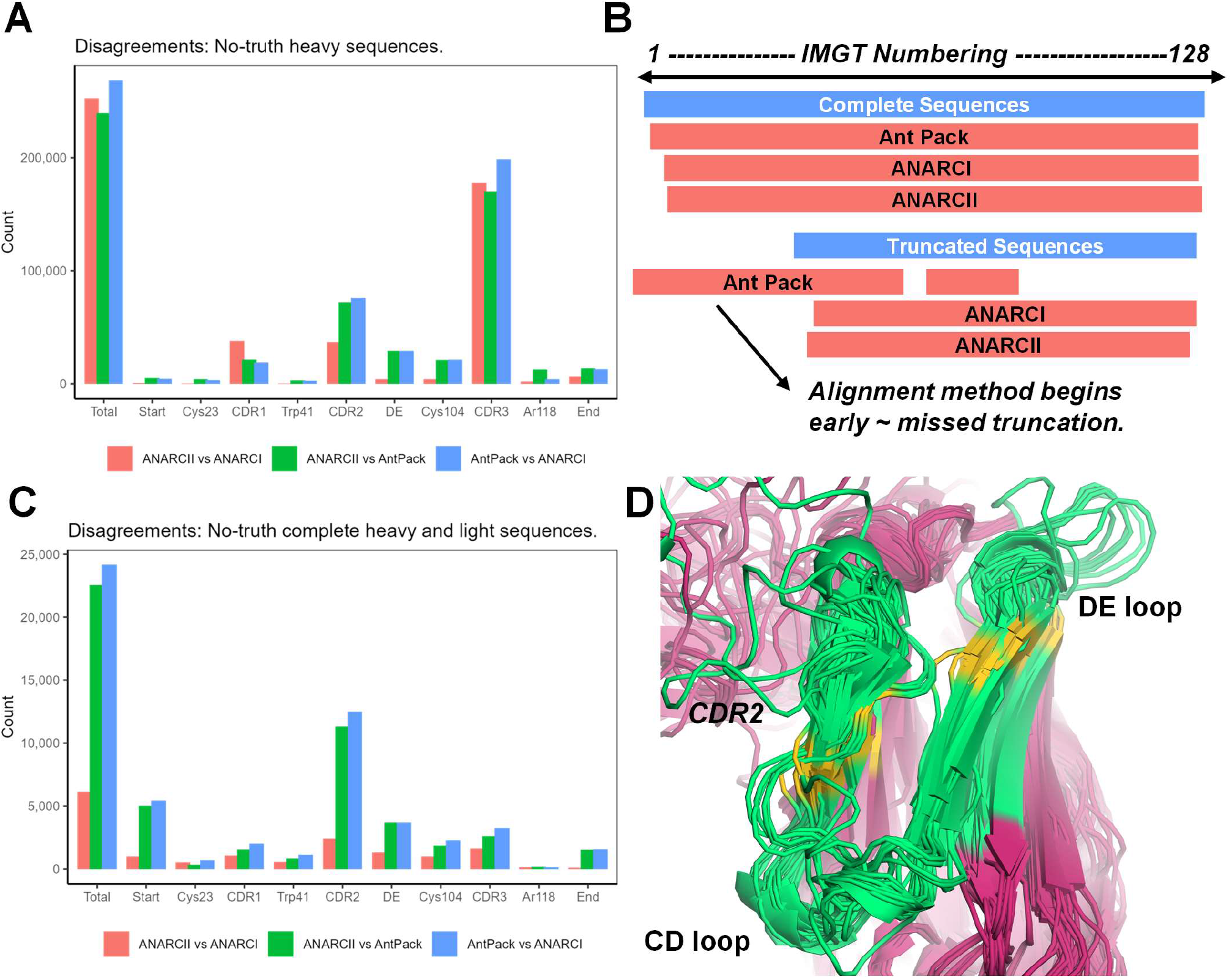
Disagreements between numbering tools highlight weaknesses of alignment methods and identify ambiguity in the CDR2-DE-loop region. (**A**) Comparison of AntPack, ANARCI, and ANARCII-accuracy on ‘no-truth’ heavy sequences (where two versions of ANARCI failed to agree) according to the sequence region where the disagreement occurs. (**B**) Schematic of how numbering tools deal with truncated sequences. (**C**) Corresponding disagreement analysis of complete heavy and light sequences found in the ‘no-truth’ set where 2 of three methods could identify the residues labelled 2 and 127. (**D**) Alignment of antibody structures in SAbDab where ANARCI and ANARCII-accuracy disagree in the CDR2 to DE loop region of the heavy chains. IMGT residues 55-90 are coloured green with from the beginning of the CDR2 to the end of the DE loop. Residues associated with the loop start/ends (CDR2: 55, 66; DE: 80, 86) are coloured in gold. PDB codes used: 3RPI, 4P9H, 4P9M, 4RWY, 4RX4, 4YDL, 5A7X, 5A8H, 5C0N, 5C7K, 5CJX, 5DMG, 5F21, 5JS9, 5JSA, 5THR, 5TRP, 5V6M, 5VIY, 5VJ6, 6CM3, 6DBD, 6EDU, 6MF7, 6NQD, 6URH, 6XJA, 6XYM, 6XZF, 6Y0E, 7F5H, 7FBI, 7JOO, 7KDE, 7MXE, 7OW1, 7OXN, 7T6X, 7UYL, 7UYM, 7YOY, 8BW0, 8C7J, 8CZZ, 8D50, 8DOK, 8G6U, 8GPI, 8GPJ, 8IM1, 8R4B, 8R4D, 8TTW.

**Figure 4.**
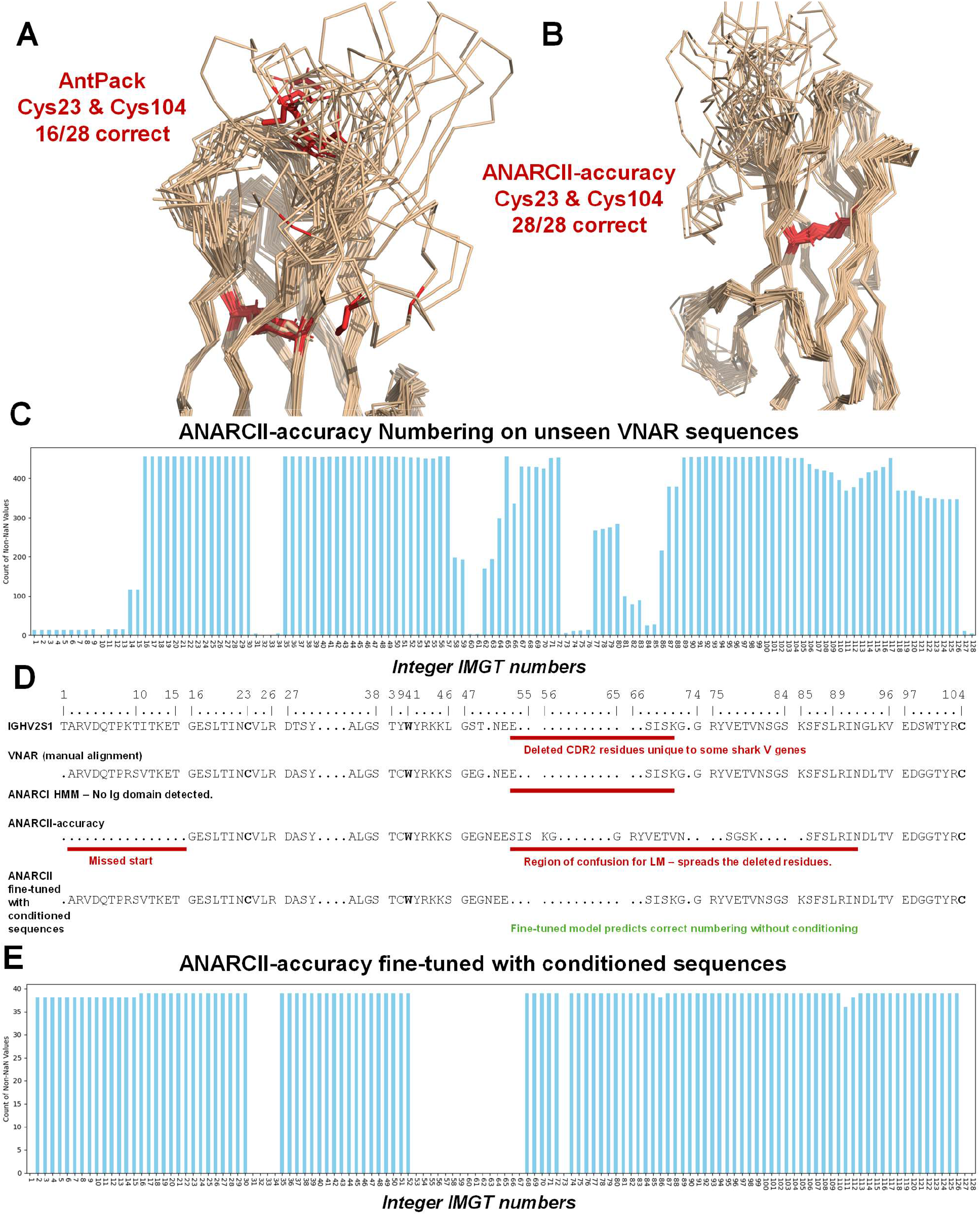
Conditioning and fine-tuning allow numbering of rare sequence types. Highlighting in red the positions identified as the conserved Cys23 and Cys104 in 28 PDB structures of VNARs numbered with AntPack (**A**) or ANARCII-accuracy (**B**). PDB codes are: 1SQ2, 1T6V, 1VER, 1VES, 2COQ, 2I24, 2I25, 2I26, 2I27, 2YWY 2YWZ, 2Z8V, 2Z8W, 3MOQ, 4HGK, 4HGM, 5L8J, 5L8K 5L8L, 6×4G, 6×4T, 7FBJ, 7FBK, 7S83, 7SPO, 7SPP, 8HGI, 8HT3. (**C**) Bar plot of IMGT number frequencies (insertions not shown) from numbering of 456 VNAR sequences by ANARCII-accuracy. (**D**) Analysis of a VNAR sequence against the closest corresponding nurse shark IMGT annotated V-gene as well as numbering by ANARCII- accuracy and a model fine-tuned on conditioned sequences. (**E**) Bar plot of IMGT number frequencies (insertions not shown) from numbering of 37 held out VNAR sequences in the test set by ANARCII-accuracy fine-tuned on conditioned sequences.

This indicated that a large fraction of discrepancies between methods were result of heavy sequence truncations or low-quality reads (presence of an ‘X’) which are commonly observed in OAS and NGS data (Olsen, Boyles, and Deane 2022). While this caused disagreements between ANARCI and *ANARCII- accuracy*, it disproportionately impacted the ability of the alignment-based method (AntPack) to correctly identify the conserved Cys23, and Cys104 positions (**Figure 3B**). For light chains in the *no-truth* data the proportion of identically numbered sequences between the three methods was much higher. Only 4.3% of sequences were not numbered identically between *ANARCII-accuracy* and AntPack, and only 1.6% between *ANARCII-accuracy* and ANARCI. Here, the proportion of mislabelled Cys23 and Cys104 by AntPack was lower (<0.1% and 0.5%), while *ANARCII-accuracy* was still the most accurate of the three methods (<0.1% Cys23/ Cys104 mislabelled) (**Supp. Fig 2D**).

Filtering for complete heavy and light sequences (where 2 of the 3 methods identified residues labelled 2 and 127), revealed that AntPack was more in line with ANARCI/*ANARCII-accuracy* and most disagreements now fell within the CDR2 to DE-loop region (**Figure 3C**). We inspected the IMGT germline numberings of this region and found that many sequence patterns in the *no-truth* set diverged substantially and had no clear way of being annotated.

We next identified structures from SAbDab where ANARCI/*ANARCII-accuracy* labels disagreed in this region to see if there was any structural pattern to aid numbering (**Figure 3D**). Our analysis (see **Supplementary Results**: ‘Structural exploration of the CDR2 to DE-loop region’) showed no consistent choice for how to number the CDR2 to DE-loop region. However, we found that both ANARCI and *ANARCII-accuracy* were consistently able to identify the conserved residues (Cys23 and Cys104) as well as agree on the CDR1 and CDR3 loops. So while the CDR2 to DE-loop region is challenging to number in some sequences it did not impact the delineation of other CDR loops.

### Identification of novel antibody sequences in the PDB missed by ANARCI

An important use case of ANARCI is to identify antibody sequences present in the PDB (Berman et al. 2000). This presented a challenge for ANARCII as the PDB contains many long sequences which exceed the length of the context window (210 tokens) and therefore require pre-processing into smaller chunks to permit identification and numbering of the antibody content. To allow the model to process long sequences we employed a combination of pattern matching and sliding window approaches to break up the sequence into fragments which are scored with a one-step decoder that identifies the region most likely to contain antibody content (**Supp Fig. 5A**, see methods). The first region predicted to contain antibody content from each long sequence is then passed to the full model for complete numbering. The model then outputs a score for each sequence derived from the logit values assigned to each integer token (non-insertion).

**Figure 5.**
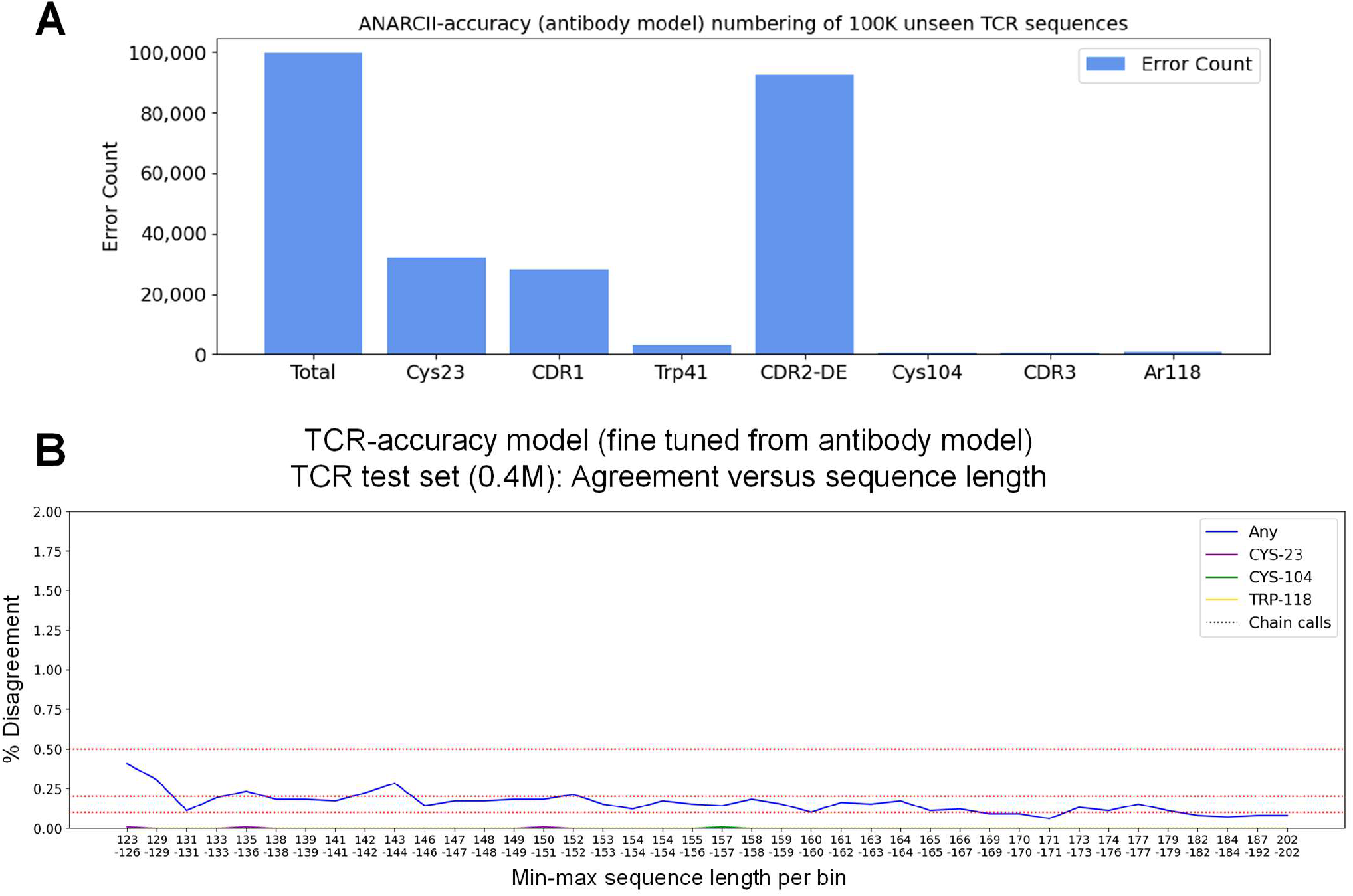
ANARCII-accuracy can identify conserved residues and CDR3 loops of unseen TCR sequences and undergo fine-tuning for complete identification. Numbering errors from ANARCII-accuracy labels on 100K TCR sequences (**A**) binned by region and conserved residues. Disagreement in numbering of ANARCI and TCR- accuracy model versus sequence length on a held-out test sets of 0.4M TCR sequences (**B**). Test sequences are divided into 40 bins based on length, indicated on the x axes, from short to longer sequences.

This method allowed us to process all sequence data from the PDB (downloaded July 2024) and demonstrated that *ANARCII-accuracy* sequence scores could be used to correctly identify over 99.9% of PDB codes previously identified as antibodies and present in SAbDab (Dunbar et al. 2014), with the model failing to identify only 3 PDB codes (engineered alanine scan variants). We then examined the distribution of sequence scores from all SAbDab data (divided by single, multi-chain or non-antibody), a set of 100K TCR sequences and a set of VHH sequences (from PLAbDab-nano (Gordon et al. 2024)) and a set of 100K UniProt sequences filtered to exclude immunoglobulins (labelled “Fails” in **Supp Fig. 5B**). *ANARCII-accuracy* could discriminate between antibodies and non-antibody proteins and assign high scores to VHH sequences despite these being subtly different from conventional (VH) heavy chain sequences.

However, in our analysis of all sequences in the PDB we found 100 PDB codes which contained antibodies and were not present in SAbDab. While most had been discounted due to issues with the structural data (missing residues in the CDRs or lack of electron density), we identified 43 sequences which were missed due to the inability of ANARCI to label them as antibodies. These sequences/structures were predominantly derived from non-human/mouse species and included rarer formats such VNARs, chimeric constructs and others simply labelled as ‘novel’ or ‘new’ immune receptors not typically recognised as conventional antibodies (for example see PDB codes: **8HT3, 3BDB, 7SPP**). In each case, the flexibility of *ANARCII- accuracy* allowed it to find a suitable numbering pattern that did not compromise identification of conserved residues and the CDR1 and CDR3 loops. This led us to explore in detail how *ANARCII-accuracy* dealt with sequences from species and types not seen in training data that standard approaches typically fail on.

### ANARCII generalises to unseen sequence types and can be conditioned to create sensible data for fine-tuning

Increasingly, researchers are exploring alternative immunoglobulin modalities to develop novel therapeutic platforms that can address disease challenges (De Pauw et al. 2023; Gauhar et al. 2021). An example are VNARs which utilise shark V-genes and contain sequence features such large gaps or enrichment of cysteines that diverge from the standard format of normal antibody heavy chains (**Supp. Fig 6**) (Cheong et al. 2020; Fernández-Quintero et al. 2022). Tools such as ANARCI and AntPack struggle to number such sequences as their methodology cannot generalise beyond the patterns captured by their reference sets/consensus sequence. When attempting to number VNARs, alignment-based methods tend to either fail to return any alignment, or else misnumber key conserved residues such as Cys104.

Figure 4 shows the numbering of 28 VNAR structures from the PDB by AntPack (**Figure 4A**) and *ANARCII- accuracy* (**Figure 4B**). In 12 of 28 sequences numbered by AntPack the Cys104 was misidentified and placed within the CDR3 (**Figure 4A**). In *ANARCII-accuracy* numbering all the Cys104 residues were placed correctly within the framework, demonstrating the model was able to generalise and produce consistent, structurally sensible numbering at key residues.

We then inspected *ANARCII-accuracy* numbering of 456 VNAR sequences retrieved from PLAbDab-nano (Gordon et al. 2024). Conserved residues Cys23, Trp41 and Cys104 were correctly numbered in 97% of sequences (444/456), with the majority of numbering beginning from position-16. However, the CDR2 deletion was not numbered consistently, with gaps being spread across residues 57 to 90 (**Figure 4C**, see example sequence in **Figure 4D**). In 278 sequences, *ANARCII-accuracy* identified an aromatic at 118, while 25 were a cysteine, with the remaining dominated by charged residues (E, K, D, H) or unlabelled (model could not confidently number beyond residue 117 for 88 sequences).

Identification of where *ANARCII-accuracy* was failing to number correctly (by comparison to IMGT germline annotations of a shark V gene, **Figure 4D**) allowed us to perform simple correction at the time of autoregressive inference, this correction is commonly referred to as “conditioning” (Ziv, Marsden, and Deane 2024). To do this we identified patterns associated with IMGT labels and overwrote the predicted token at the inference step. The corrected token was then passed to the decoder to continue numbering (see **Supplementary Methods**: ‘Conditioning steps used to number VNAR sequences’). Through application of these conditioning steps, we were able to rapidly rerun the numbering of large sets of shark sequences and obtain numbered outputs which fully aligned with IMGT germline definitions.

We then fine-tuned *ANARCII-accuracy* on these outputs to obtain a model that could correctly identify the CDR2 gap, GXGT motif and conserved residues in all cases, without any conditioning steps (**Figure 4E**). This test case demonstrates the ease of conditioning and fine-tuning to build on the fundamental sequence patterns learnt by *ANARCII-accuracy*. We have included this fine-tuned model, ‘*VNAR-accuracy’*, in the ANARCII package and webapp to allow users to explore its predictions and accurately number VNAR sequences, as well as conditioning code and instructions for customisation.

### ANARCII models are capable of numbering large portions of TCR sequences correctly, and achieve high accuracy with minimal fine-tuning

Both *ANARCII- accuracy and ANARCII-speed* models were able to number TCR sequences in a similar manner to VNARs (correct identification of conserved residues, divergence around the CDR2/DE region, **Figure 5A**), however their chain token assignments were incompatible. For comparison, providing a TCR sequence to ANARCI and restricting the model to HMMs built from antibody germlines fails to return any alignment. We were able to expand the weights to accommodate the four TCR chain types (alpha, beta, gamma, delta) and then rapidly fine-tune both models on TCR data resulting in high accuracy versions with minimal training (**Figure 5B**). We have provided options to run the *TCR-accuracy* and *TCR-speed* models within the ANARCII package and webapp.

### Alternate numbering schemes can be obtained through conversion of IMGT outputs

To number in alternate numbering schemes such as Kabat (Wu and Kabat 1970), Martin (Abhinandan and Martin 2008) or Aho (Honegger and Plückthun 2001) we have modified the methodology used by ANARCI (Dunbar and Deane 2016). Instead of converting state vectors from HMM outputs, the conversion algorithm processes the final numbered outputs from an ANARCII model and returns the updated scheme. Once a sequence, or set of sequences have been IMGT numbered the user can rapidly, and repeatedly, convert to Kabat, Martin, Chothia or Aho without having to rerun the numbering step (see user guide at https://github.com/oxpig/ANARCII/wiki). In addition, the web app provides functionality that allows users to inspect IMGT numbering alongside an alternate scheme to visualise how the new numbering diverges from IMGT labels (https://opig.stats.ox.ac.uk/webapps/sabdab-sabpred/sabpred/anarcii/). Finally, for users with complex pipelines or codebases built on ANARCI, we have provided a legacy function which can be used to convert the ANARCII outputs to the original ANARCI format to minimise disruption.

## DISCUSSION

We have trained a language model to carry out the task of antibody sequence numbering and demonstrated that it is comparable with standard approaches such as sequence alignment and hidden Markov models. Using simple augmentation of our training data, as well as assessing the score of discrete chunks of sequence, our approach was able to identify antibody sequence content from non-antibody content as well as search through long sequences to identify the regions of interest. The agreement of sequence numbering with previous approaches was extremely high. Furthermore, our model can number rare VNAR sequences where standard approaches failed to return an alignment and has also been fine-tuned to number TCR sequences.

Our final model suite allows users to number antigen receptor sequences in a fashion that is highly consistent in the face of changing reference sets and growing numbers of nonstandard query sequences. The area of antibody sequence numbering is in essence a simple task; however, the increasing quantities of diverse and esoteric/niche data require robust generalisable methods. ANARCII is designed to accelerate the work of researchers across all of the antibody and immune receptor sequence landscape.

## METHODS

### Retrieval of non-redundant heavy and light chain sequences from the Observed Antibody Space

All antibody sequences used were retrieved from the Observed Antibody Space (OAS) (Olsen, Boyles, and Deane 2022; Kovaltsuk et al. 2018). Due to the scale of data in OAS, all files were each randomly sampled to build a representative, non-redundant set of sequences. For heavy chains, a maximum of 100,000 non-redundant sequences with dataset redundancy (the number of reads detected per sequence) greater than 1 were randomly sampled from each file. To compensate for the lower quantity of light chain data, a maximum of 500,000 non-redundant sequences were randomly sampled per file with dataset redundancy of 1 allowed. Samples were combined and duplicate sequences removed. The number of sequences collected after this step were: ∼95.6 M heavy and ∼75.0 M light chains. All sequences were renumbered with ANARCI (Dunbar and Deane 2016) compiled with latest germline datasets at the time of analysis (December 2023). This process used all available data in 99.1% of OAS light chain files; however, it left many unsampled heavy sequences (4.0% of heavy files had over 100,000 sequences at redundancy greater than 1, and therefore contained sequences which had not been sampled in the first pass). These unsampled sequences were then used to increase the numbers of rare insertions and non-human or mouse species contained in the dataset.

### Enrichment of heavy chain sequences for rare insertions and species

We analysed the numbering of all sequences in the heavy dataset and compiled statistics on the number of times each insertion was seen, for example 112A was seen over 57,448,967 times, whereas 12A was seen only 39 times. We identified all insertion codes that were seen less than 1000 times and resampled all files in OAS that were not fully sampled by the initial processing steps to find sequences with these rarer insertions and added these to the heavy dataset. After this step we enriched for non-human and non-mouse species; these included rat, rabbit, camel and rhesus macaques. Files from these species which were partially sampled from previous steps were further sampled to search for any sequences not already present in the heavy dataset. These steps added 123,305 sequences to the dataset and resulted in a final number of ∼95.7 M heavy chains. The total number of unique number labels rose from 925 pre-enrichment to 968 post-enrichment.

### Sequence clustering and dataset splits

The levels of dataset diversity in the light and heavy chain are very different, for example only 143,563 light CDR3 sequences contained insertions at position 112A (the first CDR3 germline insertion position in IMGT numbering) in the ∼75.0 M sequences, compared to ∼57.1 M in ∼95.7 M for the heavy chain. As a result of this we used different parameters for light and heavy sequence clustering to create large numbers of diverse clusters that could be split into train, test and validation sets. Using CD-Hit (Li and Godzik 2006) heavy sequences were clustered at 75% identity, while light sequences (kappa and lambda separated) were clustered at 85% identity across the entire sequence. CD-Hit parameters (aS=0.9 for light chains, aS=0.8 for heavy) were set to cluster sequences in such a way that short fragments of a longer sequence were always included in the same cluster as the parent and thus truncations of the same sequence were not present in train and test or validation datasets. This resulted in 1,067,916 heavy clusters, 1,246,933 lambda clusters and 1,837,998 kappa clusters which were each split 90:5:5 into train, validation and test sets. The final numbers of clusters and sequences in each set are shown in **Supp Table 1**.

### TCR dataset generation

Paired TCR sequences were extracted from the Observed TCR Space (OTS) (Raybould et al. 2024), and unpaired sequences from iReceptor (Corrie et al. 2018) (for details see Raybould et al. 2024, raw data can be found at https://doi.org/10.5281/zenodo.11208211). Paired sequences were split into individual chains, combined with unpaired data and numbered with the latest version of ANARCI. As ANARCI can falsely identify a certain subset of alpha chains sequences with TRAV/DV genes as delta chains, we utilised IgBLAST (Ye et al. 2013) genes calls where available. Using this process 435 chains in the training data were delta chains. In total 4,993,877 alpha, 6,067,942 beta and 526 gamma chains were collected for training TCR-specific models.

### Addition of junk and tokenisation of training data

For each sequence in the train and validation datasets between 0 and 40 random amino acids were added either side of the antibody content to mimic natural non-antibody content often found in antibody sequence datasets and crystal structures. The corresponding target labels of these randomly added residues were defined by a [<SKIP>] token. Starting junk residues were added after the [<SOS>] token and before true antibody content began. After antibody content ended, an [<EOS>] token was added, followed by random junk residues. The purpose of this was to a provide an easy way to identify the end of the sequence during post inference processing of predicted tokens. The amino acid tokeniser also includes a token for non-standard residues (e.g. B, O, J, U, X, Z) [X] which may represent unknown residues in sequencing methods like NGS or mass spectrometry. The numbering tokeniser used to create target labels included all numbers from 1-128, as well as an [X] to mark an insertion. All insertion labels were collapsed to [X] and are translated to the correct letter codes in post processing. The input and output tokens pass through different embedding layers in the encoder and decoder respectively, therefore shared tokens like [X], [H], [L], [K] each have separate learned representations within the source and target sequences.

### Model Training

Due to the large amount of training data (∼140M sequences, comprised of ∼70.5M heavy, ∼38.7M kappa and ∼29.9M lambda chains), heavy and light sequences were each split into 5 subsets, of which ∼10M heavy sequences, ∼6M kappa sequences and ∼4M lambda were randomly sampled and passed to a dataloader. Validation loss was calculated after all 5 heavy and 5 kappa/lambda subsets had been randomly sampled (in random order) meaning that the model had seen exactly 50M heavy and 50M light datapoints per epoch by our definition. Validation loss was calculated separately for heavy and light sets, then the average validation loss across both chains used for final model selection. Batch size of 512 was found to provide optimal validation loss during testing. Models design and training was performed in PyTorch version 2.2.1 (Paszke et al. 2019), Cuda version 12.1 and Python version 3.11. Model validation and testing was performed in multiple versions of Python (3.9-3.13), PyTorch (2.0-2.5) and Cuda (11.8, 12.1, 12.4) to ensure performance was consistent, and run parameters and calls were all functional across platforms when called from the final package.

### Model architecture and hyperparameter selection

We utilised the standard Seq2Seq architecture described by Vaswani et al. (Vaswani et al. 2017). A set of model hyperparameters were chosen based on convergence over 15-30 epochs (full details shown in **Supp Table 2**, along with corresponding inference speeds on 10,000 sequences). Several models were carried forward to longer training cycles, with the balance of validation accuracy versus inference speed being used to settle on two sets of hyperparameters: a 1-layer model (encoder/decoder hidden dims of 128) with slight reduction in accuracy but faster inference speed (*ANARCII-speed*), and a 2-layer model (also 128 hidden dims, **Supp Figure 1**) which was slower but had higher accuracy (*ANARCII-accuracy*). Ever larger models did give marginal increases in accuracy but also trebled the inference speed relative to the smallest model (**Supp Table 2**).

### Training regime

Three training regimes were tested: a standard step decay learning rate, cosine decay with warm restarts and a warmup phase with cosine decay. Cosine decay with warm restarts every 15 epochs (initial learning rate = 0.0002, minimum = 0.00001) resulted in models with the best validation loss. Each model (*ANARCII-speed* and *ANARCII-accuracy*) was trained in this way on an Nvidia-A100-GPU for 75 epochs, then trained for a further 60 epochs with initial and minimum learning rates reduced by a factor of 10 and restarts every 10 epochs.

### Processing of long sequences

The ANARCII models utilised an encoder with a maximum input length of 210 tokens, which therefore could not process very long sequences which are often found in the PDB (Berman et al. 2000). To permit ANARCII models to number the antibody content of longer sequences we utilised two strategies to find the relevant 200 residue region which could be passed to the model to output full numbering. Firstly, for each sequence over 200 residues we perform a compiled regex search for instances of C, W, C separated by 5-25 and 50-80 residues respectively. For every match we pass the content between the two cysteines to the ANARCII model and a perform a rapid, single forward pass through the decoder to obtain a score per match derived from the logit value of the predicted chain token (analogous to a <CLS> token). The first high scoring match is selected, and upstream/downstream sequence added from the beginning cysteine (−40, +160), to give a final sequence length of 200 which is fed back into the full model for complete numbering. If no CWC matches are found, or no high-scoring match is found that exceeds a defined threshold then a second strategy is employed. Here, the long sequence is broken up into sliding windows of 90 residues (along an increment of 3 residues) and each window run through ANARCII in a single forward pass of the decoder. The first high scoring window including flanking sequence (−40 from start, +70 from end, giving 200 residues) is then passed into the full model for complete numbering.

### Creation of the ‘*no-truth*’ dataset

All data collected for antibody heavy and light chains was retrieved from OAS with the ANARCI numbering created at the time at which the datasets were first processed (between 2018-2022). Each sequence was also renumbered with a more recent version of ANARCI (compiled December 2023) and the two numbering versions compared. This highlighted ∼3.6M heavy (3.8% of total heavy) and ∼0.9M light sequences (1.2% of total light) where there was some disagreement between ANARCI versions. We filtered these to find sequences with large variation in number labels (over 4 different number labels between the two versions) resulting in 530,804 heavy and 410,148 light sequences that we refer to henceforth as the ‘*no-truth*’ dataset.

### Conditioning of outputs for numbering of VNAR sequences

ANARCII-accuracy was used to number 456 shark VNAR sequences retrieved from PLAbDab-nano (Gordon et al. 2024). These numberings were compared to the IMGT germline reference for nurse shark V-genes (those with a large CDR2 deletion e.g. IGHV2S1) to understand where the language model predictions differed. The autoregressive inference step was modified to search for sequence patterns and predicted numbering (using regular expressions) that corresponded to where the predictions differed from the germline reference set. When one of these patterns was detected, the number labels were changed to exactly match the IMGT reference labels. The corrected numbering was then fed back into the decoder to continue next token prediction of number labels for the remaining sequence until completion. A detailed description of the sequence patterns and corresponding numbering can be found in the supplement (see **Supplementary Methods**: ‘Conditioning steps used to number VNAR sequences’). A Jupyter notebook containing all code and instructions on how to perform and customise the conditioning of number labels can be found at: https://github.com/oxpig/ANARCII/blob/main/notebook/conditioning.ipynb.

### Fine-tuning on shark VNAR sequences numbered by conditioning

VNAR sequences which had been numbered through conditioning of *ANARCII-accuracy* outputs (see previous section) in which all conserved residues (Cys23, Cys104 and Trp41) were correctly identified and contained the GXGT motif marking the end of the CDR3 (397 out of 456) were randomly split 80:10:10 into train, validation and test sets. Datasets were up sampled by a factor or 10, randomly shuffled and junk sequence added (see previous section, ‘Addition of junk sequence and tokenisation of training data’). The *ANARCII-accuracy* model was fine-tuned on these data for 80 epochs at a reduced batch size of 256 (warm restarts every 10 epochs, initial learning rate = 0.0002, minimum = 0.00001).

### Fine-tuning on TCR data

TCR sequences (see TCR dataset generation section) were clustered using CD-Hit at the level of 85% identity across the entire sequence and split 90:5:5 into train, test and validation sets. Sequences were numbered with the most recent version of ANARCI (compiled December 2023). Junk sequence was randomly added to the start and end of each sequence as was carried out for antibody sequences (see previous section, ‘Addition of junk sequence and tokenisation of training data’). Fine-tuning was performed on both antibody-specific models (*ANARCII-accuracy* and *ANARCII-speed*) with weights expanded to accommodate the TCR specific tokens for chain call (A, B, D, G). Models were fine tuned for 80 epochs using identical learning rate parameters to the second stage of training at a reduced batch size of 256 (warm restarts every 10 epochs, initial learning rate = 0.0002, minimum = 0.00001).

### Sequence classifier for unknown sequences

A classifier was developed to facilitate a two-step “unknown” mode where input sequences are first classified as either antibody or TCR, and then passed to the correct numbering model. The classifier model architecture was identical to the *ANARCI-speed* model (1-layer, 4 heads and 128 hidden dims in the encoder and decoder), only differing in the decoder output where a single chain token was predicted (<TCR> or <Ig>). The model was trained using sequences sampled from the train, validation and test datasets generated previously (see sections, ‘*Sequence clustering and data splits*’ and ‘*TCR dataset generation*’). For the antibody data: heavy, kappa and lambda sequences were each separately sampled to obtain an even distribution across the clusters found using CDHit (W. Li and Godzik 2006). This resulted in ∼3.9M heavy, ∼4.2M kappa and ∼4.7M lambda training sequences, with ∼0.3-0.8M sequences in the corresponding test and validation sets. All TCR sequences from train, validation and test sets were used resulting in ∼4.4M alpha and ∼5.5M beta training sequences with ∼0.3-0.4M sequences in the corresponding test and validation sets (gamma and delta sequences were highly underrepresented as previously described). The model was trained for 80 epochs using cosine decay (learning rate parameters identical to initial stage of training for ANARCII-speed, batch size of 256), before reaching 100% accuracy on the validation set.

### Webapp and package

The suite of ANARCII language models can be selected and run from a webapp (https://opig.stats.ox.ac.uk/webapps/sabdab-sabpred/sabpred/anarcii) where output files can be downloaded in csv or MessagePack (msgpack) format. Users can visualise the differences between IMGT numbering and the alternate scheme requested through a side-by-side sequence analysis tool. The suite is also available to download (https://github.com/oxpig/ANARCII).

## CONTRIBUTIONS

CMD, SAR, AGW and PA conceptualised the study and designed the methodology. SAR, AGW and PA curated data used to train models. AGW, SAR and PA trained models. AGW, GLG and FCS curated extra exploratory data. AGW, BW, BAS, and FCB developed software and web tools. AGW, PA, YL, HLC, GLG and CMD analysed the data. AGW visualised the data. CMD provided computational resources. AGW wrote the original manuscript draft. CMD reviewed/edited the manuscript and supervised the project.

## AI DISCLOSURE

Generative AI tools, including ChatGPT and GitHub Copilot, were utilised to assist in code generation and error checking during the development of this project.

## ACKNOWLEDGEMENTS

The authors would like to thank Oliver Turnbull, Carlos Outeiral and David Prihoda for their helpful suggestions and feedback.

