## Supplementary Information for "ANARCII: A Generalised Language Model for Antigen Receptor Numbering"

### SUPPLEMENTARY METHODS

**Conditioning steps used to number VNAR sequences.** Through inspection of *ANARCII-accuracy* numbering on VNAR sequences from PLabDab-Nano (Gordon et al. 2024) we identified key patterns which preceded 'misnumbering' by the language model (according to IMGT germline annotations of a shark V gene, **Figure 5D**). These patterns were used to perform targeted intervention in the autoregressive inference loop and overwrite the predicted token with the correct token, and then pass the partial sequence into the decoder to continue the inference loop.

- To correct the misidentification of the starting sequence we identified the beginning pattern being missed and labelled the first residue according to IMGT numbering (often 1 or 2).
- Next, we used regular expressions to identify the sequence associated with the deleted CDRH2 region (often flanked by NEE) and labelled the subsequent sequence starting from residue 68 to preserve the large deletion characteristic of VNARs.
- Finally, the CDRH3 was completed (often the model would stop numbering long CDRH3s) by identifying the conserved cysteine at position 104 and the terminal GXGT motif at positions 119-122. Once these two patterns were found then the intermediate sequence was renumbered according to standard rules. Through application of these conditioning steps, we were able to rapidly rerun the numbering of large sets of shark sequences and obtain "correctly" numbered outputs.

Finally, we took the conditioned sequences and filtered to obtain those with conserved cysteines at 23 and 104, tryptophan at position 41 as well as the presence of a GXGT motif at positions 119-121 resulting in 397 complete sequences. We split this dataset (80:10:10) and used it to rapidly fine tune the *ANARCII-accuracy* model. When challenged with 38 sequences from the held-out test set, the fine-tuned model was able to correctly identify the CDRH2 gap and GXGT motif and conserved residues in all cases, and the starting number in 37 of 38 sequence (**Figure 5E**).

### SUPPLEMENTARY RESULTS

**Structural exploration of the CDR2 to DE-loop region in sequences where numberings differ between ANARCII and ANARCI.** To systematically investigate how numbering aligned with secondary structural features such as loops or beta sheets, we used DSSP (Touw et al. 2015) to analyse all SAbDab structures numbered with either ANARCI (**Supp Figure 4A**) or *ANARCII-accuracy* (**Supp Figure 4B**). This revealed a distinct secondary structure pattern alternating between extended strands (coloured pink) and turns/bends (yellow/sand) that were synchronised with IMGT numbering. We isolated 113 SAbDab chains (57 PDB codes) in which ANARCI and *ANARCII-accuracy* disagreed at the CDR2 to DE-loop (but agreed on the preceding conserved Cys23 and Trp41 residues). Visualising the IMGT numbering-to-structure relationship with DSSP revealed a complex pattern with many insertions, underlining the ambiguity in numbering this region (**Supp Figure 4C-D**). This could also be seen structurally from alignment and inspection of the 3D coordinates of the heavy chains (**Figure 3D**) with a large variation in the length and conformation of the DE loop, the distal CD loop and small variations in the bridging framework regions. For both tools, numbering mostly aligned with structural features, with differences arising in placement of insertions and deletions after the CDR2 and either side of the DE-loop (**Supp Figure 4C-D**).

### SUPPLEMENTARY FIGURES

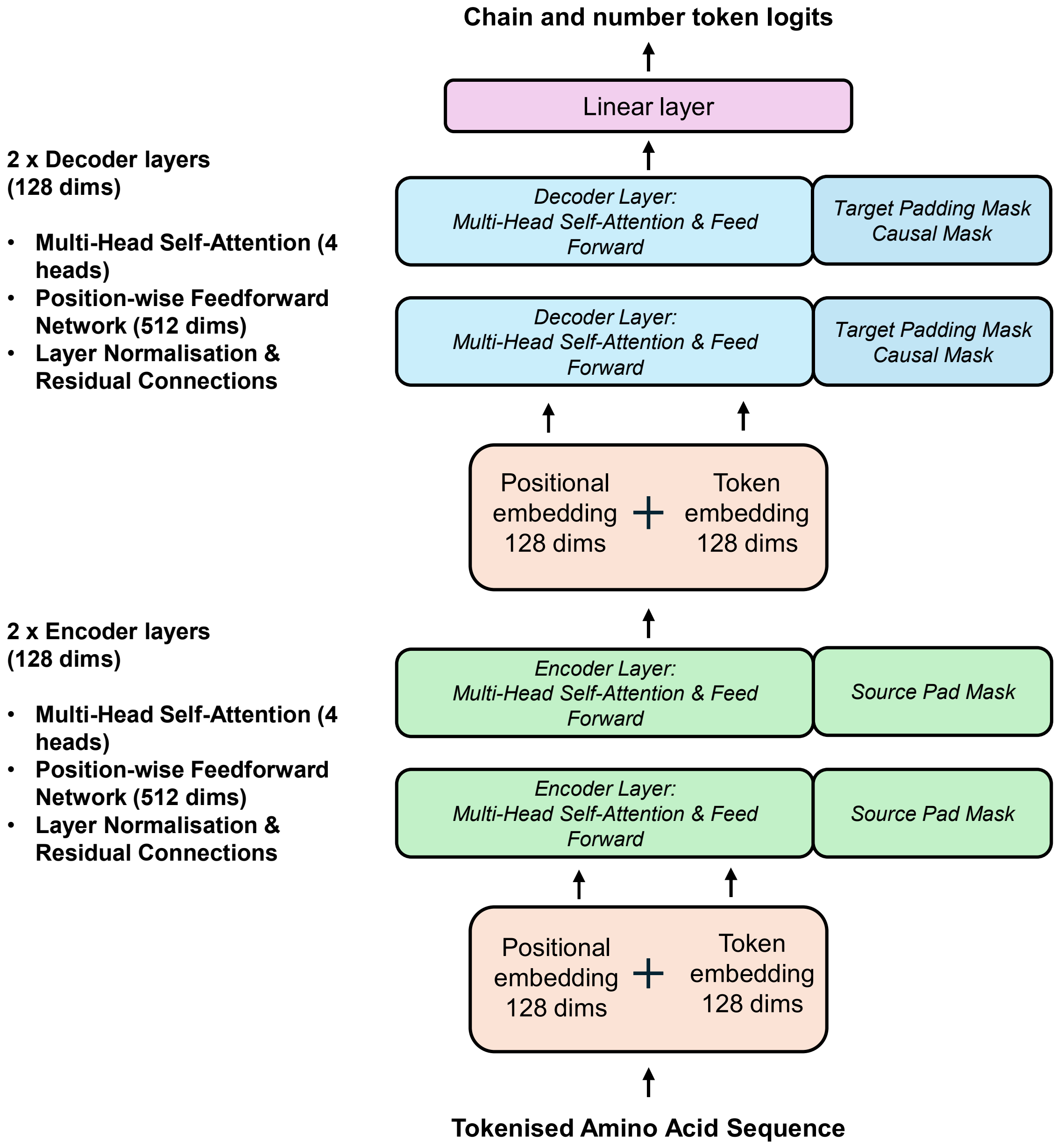


***Supplementary Figure 1: Model architecture of ANARCII-accuracy.*** *Schematic detailing the Seq2seq architecture of ANARCI—accuracy (ANARCII-speed has identical dimensions with only 1 encoder and 1 decoder layer). Blocks indicate the components of the encoder-decoder transformer model used to assign chain and number token logits to input amino acid sequences. The input is a tokenised amino acid sequence (tokens include 20 standard residues, nonstandard residues: X, delimiters: SOS, EOS, PAD), which is first embedded using token and positional embeddings (128 dimensions each), summed and scaled. This representation is passed through 2 encoder layers (128 dimensions, 4 attention heads), each consisting of multi-head self-attention, position-wise feedforward networks (512 dimensions), layer normalisation, and residual connections. The encoded input sequence is then passed to the decoder, which processes the representation using the same architecture (2 decoder layers, 128 dimensions, 4 attention heads) and additional causal and padding masks. The final decoder output is projected through a linear layer to 136-dimensional logits, corresponding to chain (H,K,L), insertion (X), delimiter (SOS, EOS, SKIP, PAD) and number label (1-128) tokens.*

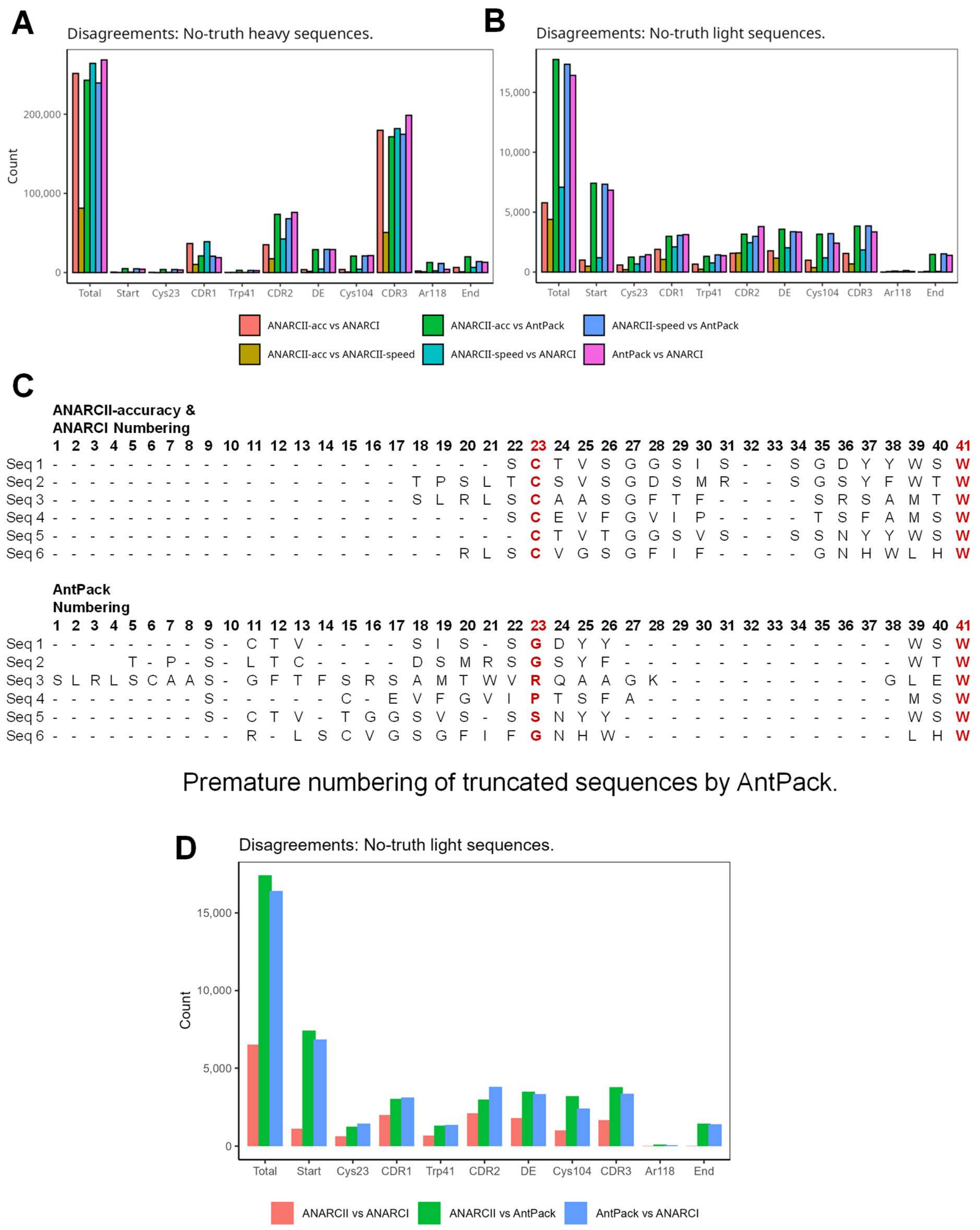


***Supplementary Figure 2: Analysis of disagreements between numbering tools on ambiguous sequences.*** *Comparison of AntPack, ANARCI, ANARCII-accuracy and ANARCII-speed on 'no-truth' heavy (**A**) and light (**B**) sequences (where two versions of ANARCI failed to agree) according to the sequence region where the disagreement occurs. (**C**) Examples of ANARCI/ANARCII-accuracy versus AntPack numbering of truncated sequences corresponding to schematic in Figure 3B. Specifically detailing the early start by AntPack which misses the truncation identified by ANARCI and ANARCII-accuracy. (**D**) Comparison of AntPack, ANARCI, and ANARCII-accuracy on 'no-truth' light sequences (identical to data in part B, however with less comparisons to allow for better readability).*

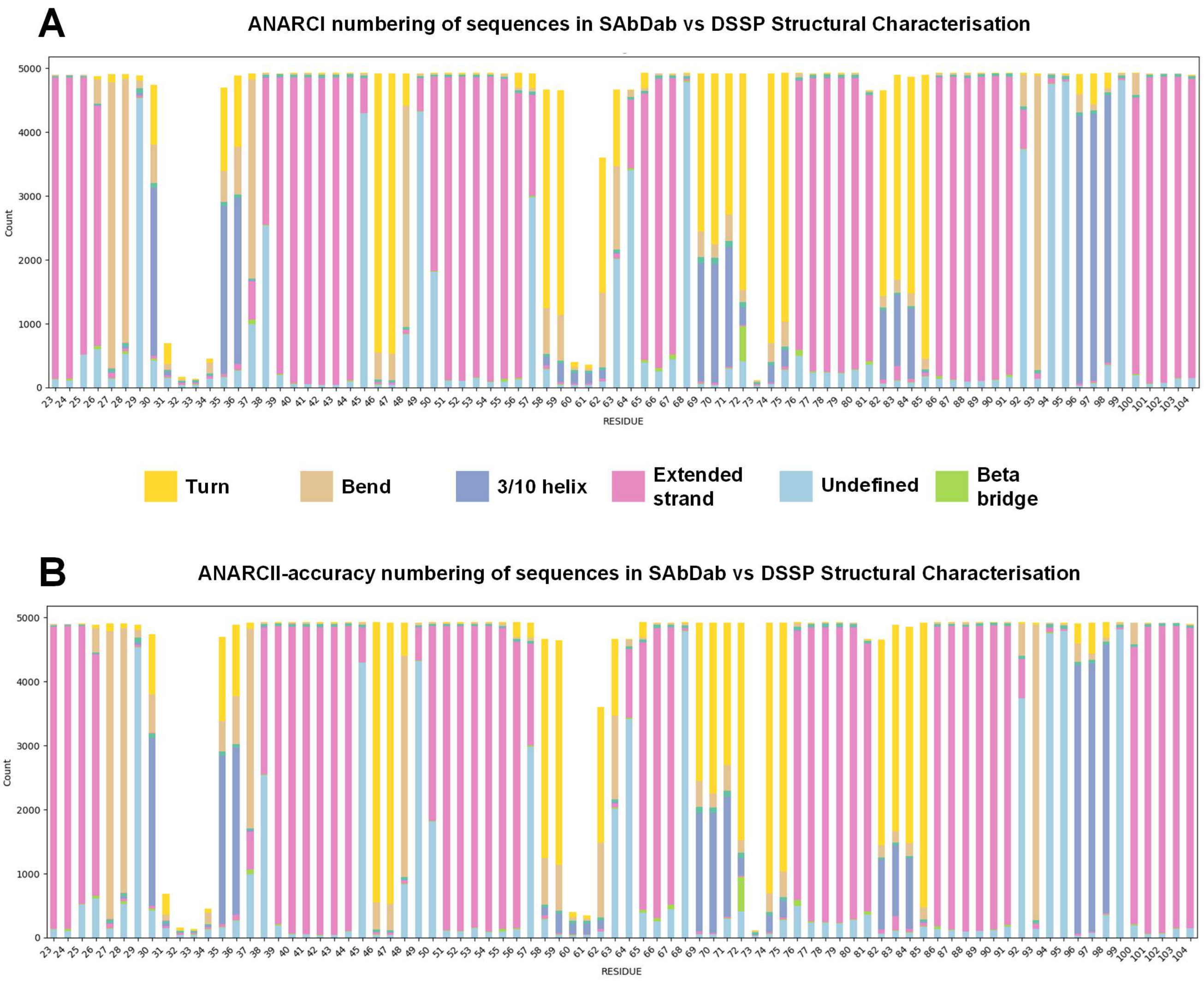


***Supplementary Figure 3: Structural analysis of ANARCI and ANARCII-accuracy numbering of sequences in SAbDab.*** *Dictionary of protein secondary structure (DSSP) characterisation of SAbDab structures numbered with ANARCI (**A**) or ANARCII-accuracy (**B**). Bar plots show the proportion of DSSP classes by IMGT integer position (insertions not shown) in heavy chains for X-ray crystal structures found in SAbDab (one heavy chain analysed for 4945 PDB codes). Clear patterns of secondary structure associated with loops and frameworks align to the assigned IMGT numbering scheme.*

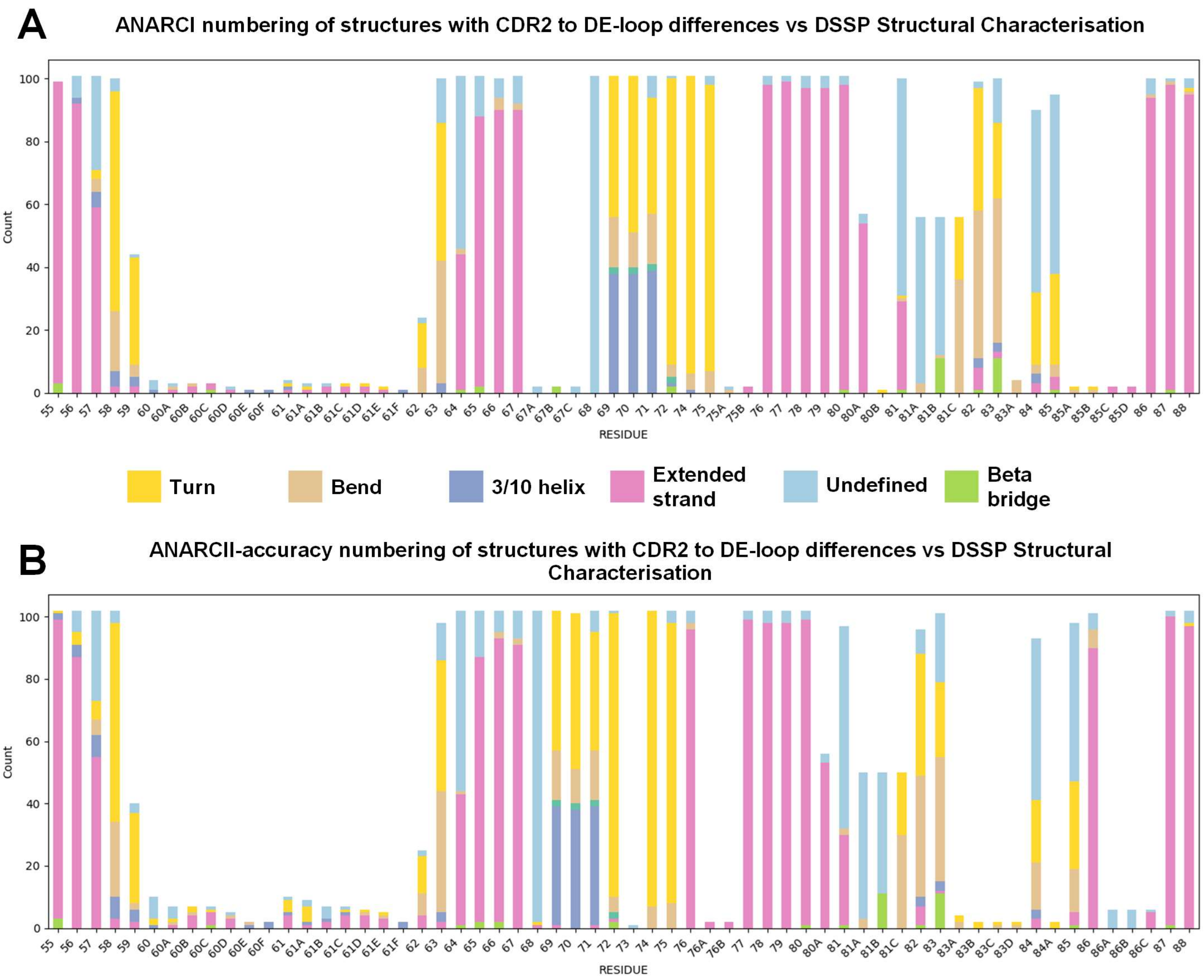


***Supplementary Figure 4: Structural analysis of ANARCI and ANARCII-accuracy numbering of sequences in SAbDab which differ in the CDR2 to DE-loop region.*** *Dictionary of protein secondary structure (DSSP) characterisation of SAbDab heavy chains where the numbering of the CDR2 to DE-loop region disagreed between ANARCI (A) and ANARCII-accuracy (B), corresponding to 113 chains from 57 PDB codes. Bar plots show the proportion of DSSP classes by IMGT position including insertions.*

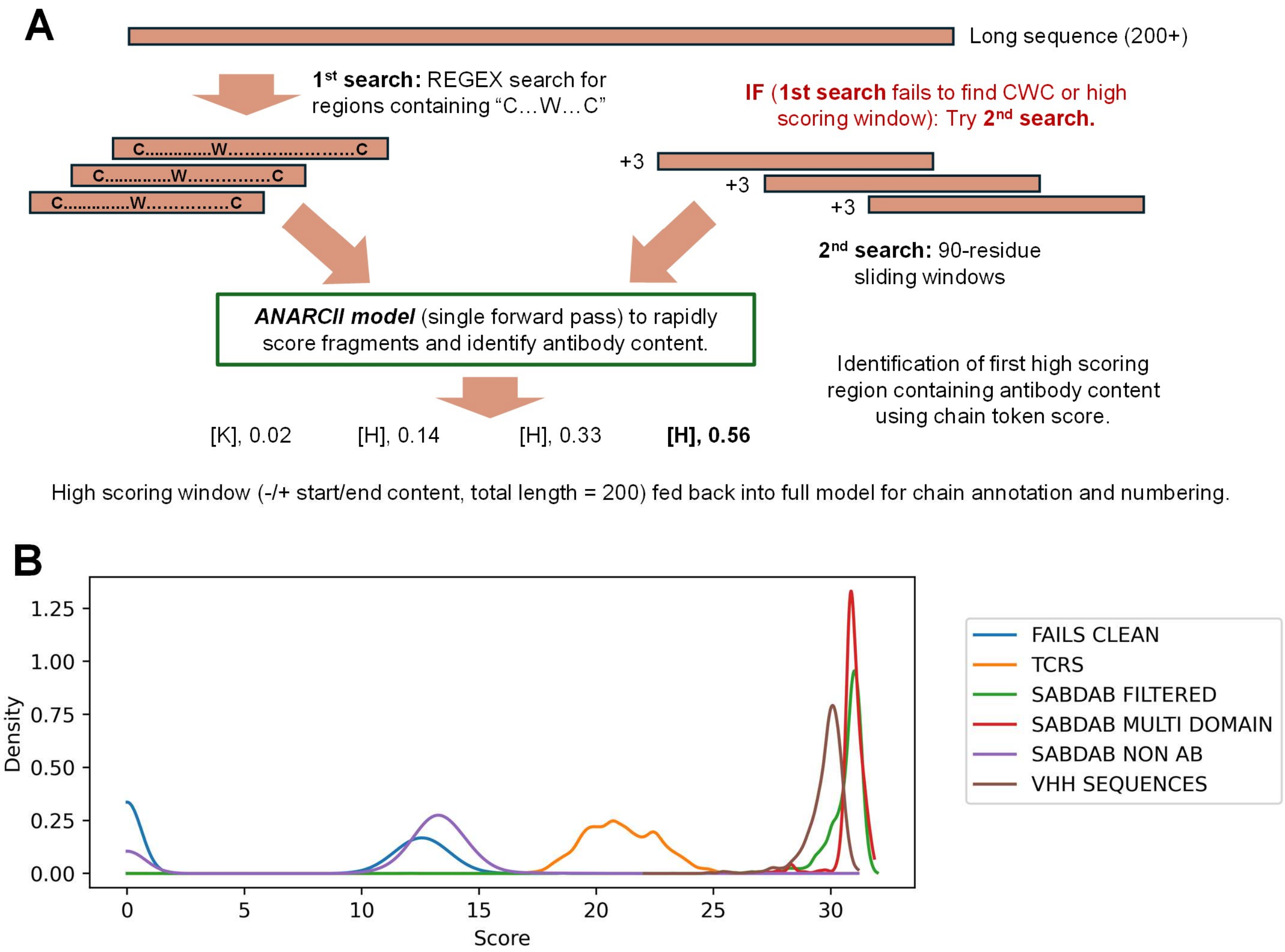


***Supplementary Figure 5: Preprocessing steps allow numbering of long sequences present in SAbDab.*** *(**A**) Sequences longer than 200 amino acids are first screened for a CWC pattern to find chunks which may represent antibody content. The regions with CWC matches are each passed to ANARCII in a single forward pass of the decoder to obtain the corresponding score of the chain token. The first match which scores over a defined score threshold is then fed back into the full model with extra sequence at the start (-40 from the primary cysteine) and end (+160 from the primary cysteine) to ensure complete numbering. If no window passes the threshold or no CWC matches are found, then each sequence is broken up in 90 residue windows which increment every 3 residues. These windows are then scored as before, with the first window over a threshold being run back into the model (-40, window, +70). If no window passes the threshold, then the highest scoring window is chosen. Density plot of sequence scores output by the model (mean of top token scores) for numbered sequences derived from SAbDab, 100,000 TCR sequences, PLAbDab-nano (VHH) and 100,000 non-antibody sequences (Fails) retrieved from UniProt (**B**). SAbDab sequences are divided into non-antibody chains (NON AB), multi-domain chains containing more than one FV regions (MULTI DOMAIN) and chains with only on FV region (FILTERED).*

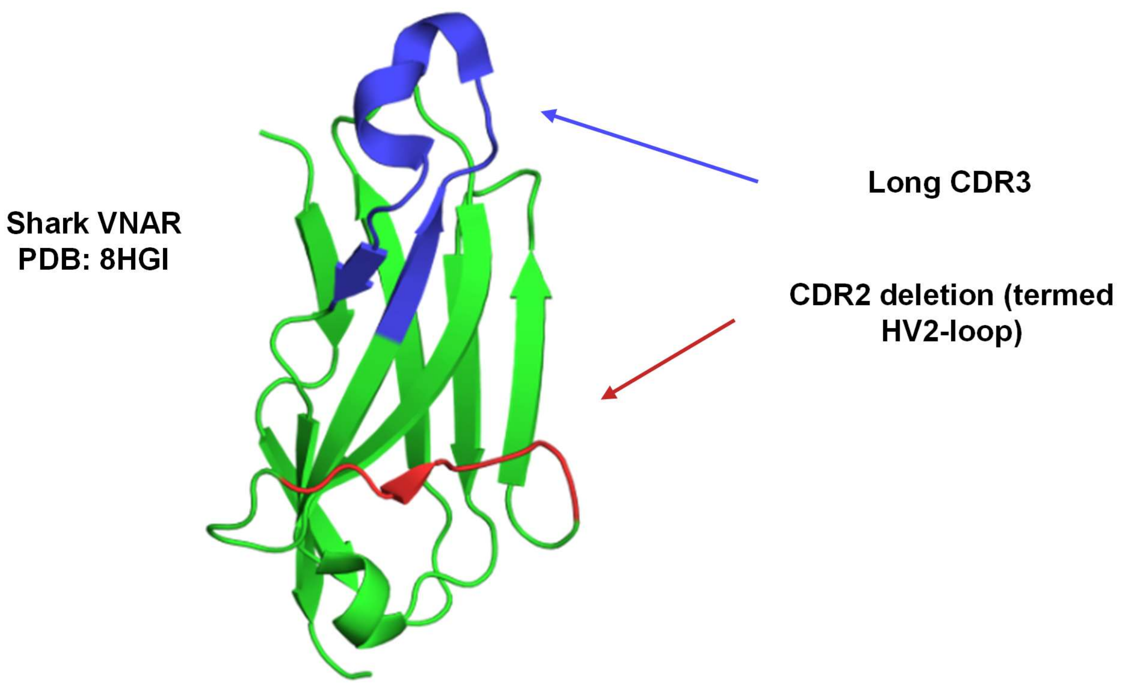


***Supplementary Figure 6: Structural features that characterise Shark VNAR sequences.*** *Example of structural features of shark VNARs which diverge from standard antibody heavy chains. The deletion at the CDR2 (HV2-loop) is shown in red, and long CDR3 shown in blue.*

### SUPPLEMENTARY TABLES

| | Train Clusters | Train Seqs | Validation Clusters | Validation Seqs | Test Clusters | Test Seqs |
|---|---|---|---|---|---|---|
| **Heavy** | 961,124 | 73,762,241 | 53,396 | 2,723,809 | 53,396 | 19,258,862<br><br>(random split into 2 subsets) |
| **Kappa** | 1,654,198 | 38,715,223 | 91,900 | 1,371,630 | 91,900 | 1,527,549 |
| **Lambda** | 1,112,239 | 29,922,215 | 62,347 | 1,100,496 | 62,347 | 2,359,102 |

***Supplementary Table 1: Numbers of clusters and sequences.***

| Name | Layers Enc/Dec | Hidden Dims Enc/Dec | Position-wise FF Dims Enc/Dec | Heads | Validation loss after 15 epochs | Best validation loss after 60-100 Epochs | Size (MB) | ~Seqs per minute (A100- without optimisation) |
|---|---|---|---|---|---|---|---|---|
| **Speed** | 1 | 128 | 512 | 4 | 0.000860 | 0.000797 | 2.2 | 55,000 |
| **Accuracy** | 2 | 128 | 512 | 4 | 0.000776 | 0.000732 | 4.0 | 34,000 |
| **Super1** | 1 | 256 | 1024 | 4 | 0.000791 | 0.000748 | 7.9 | 30,960 |
| **Super2** | 2 | 256 | 1024 | 4 | 0.000750 | 0.000716 | 15.0 | 18,969 |

***Supplementary Table 2: Model hyperparameters and inference speed (without later optimisations in data processing and value-caching to improve speed).***
